## Supplementary material for "Human genetic ancestry, *Mycobacterium tuberculosis* diversity and tuberculosis disease severity in Dar es Salaam, Tanzania": SuplementaryFiles

**Supplementary Table S1 - The different ethnic groups with at least 10 members in our cohort and the region and broad geographic location of the original area of the ethinic group.** The latitude and longitude are given in decimal degrees. In the case of two associated regions, a location close to the border of the two regions was selected.

| Ethnic group | Region | Location | Latitude | Longitude |
|---|---|---|---|---|
| Rangi | Dodoma | central | -6.5738 | 36.2631 |
| Digo | Tanga | northeast | -5.06667 | 39.1 |
| Gogo | Dodoma | central | -6.5738 | 36.2631 |
| Hehe | Iringa | southcentral | -7.7681 | 35.6861 |
| Jita | Mara | northern | -1.25 | 34.15 |
| Makonde | unclear | southeast | - | - |
| Nyakyusa | Mbeya | southern | -8.9094 | 33.4608 |
| Nyamwezi | multiple | westcentral | - | - |
| Sukuma | African Great Lakes | northwest | - | - |
| Zaramo | Dar es Salaam and Pwani | eastcentral | -7.3238 | 38.8205 |
| Chaga | Kilimanjaro | northeast | -4.1337 | 37.8088 |
| Haya | Kagera | northwest | -1.91667 | 31.3 |
| Kwere | Pwani | eastcentral | -6.3368 | 38.3939 |
| Ndengereko | Pwani | eastcentral | -7.3238 | 38.8205 |
| Ngindo | multiple | southeast | - | - |
| Pare | Kilimanjaro | northeast | -4.2678 | 37.9315 |
| Pogolo | multiple | central | - | - |
| Shambaa | Tanga | northeast | -4.75 | 38.5 |
| Yao | Ruvuma and Mtwara | southern | -10.922551 | 38.00335 |

| | | | | |
|---|---|---|---|---|
| Zigula | northern Pwani, southern Tanga | northeast | -5.77759 | 37.81088 |
| Bondei | Tanga | northeast | -5.42254 | 38.96151 |
| Ha | Kigoma | northwest | -4.8824 | 29.6615 |
| Luguru | Pwani and Morogoro | eastcentral | -7.21488 | 38.35386 |
| Makua | Mtwara | southern | -10.6417 | 39.2376 |
| Matumbi | Lindi | southern | -9.1497 | 38.9877 |
| Mwera | Kilwa district | southern | -9.1497 | 38.9877 |
| Ngoni | multiple | southern | - | - |
| Nyaturu | Singida | northcentral | -6.7453 | 34.1532 |
| Shirazi | Swahili coast | eastern | -5.72573 | 39.29856 |
| Ware | Mara | northern | -1.7754 | 34.1532 |

**Supplementary Table S2 - Phylogenetic markers selected to identify the Introductions.** The position is based on the reconstructed reference of the ancestor (79) and the derived base indicates the base present in the respective Introduction. Intro1 refers to Introduction 1 within L2.2.1, Intro5 to Introduction 5 within L4.3.4, Intro9 to Introduction 9 within L1.1.2, and Intro10 to Introduction 10 within L3.1.1.

| Lineage | Position | Derived base | Introduction |
|---|---|---|---|
| L4 | 1159734 | A | Introduction 5 |
| L4 | 1502120 | A | Introduction 5 |
| L4 | 2090889 | A | Introduction 5 |
| L4 | 225495 | C | Introduction 5 |
| L4 | 2484751 | G | Introduction 5 |
| L4 | 2624654 | T | Introduction 5 |
| L4 | 3840719 | C | Introduction 5 |
| L2 | 119555 | T | Introduction 1 |
| L2 | 1252462 | G | Introduction 1 |
| L2 | 128469 | A | Introduction 1 |
| L2 | 1547909 | A | Introduction 1 |
| L2 | 1608967 | G | Introduction 1 |
| L2 | 2111284 | A | Introduction 1 |
| L1 | 1348760 | T | Introduction 9 |
| L1 | 1392225 | T | Introduction 9 |
| L1 | 1459473 | T | Introduction 9 |
| L1 | 1479463 | G | Introduction 9 |
| L1 | 1578635 | T | Introduction 9 |
| L1 | 1648566 | A | Introduction 9 |
| L1 | 1750150 | T | Introduction 9 |
| L1 | 1956763 | C | Introduction 9 |

| | | | |
|---|---|---|---|
| L1 | 1968814 | T | Introduction 9 |
| L1 | 2189095 | T | Introduction 9 |
| L3 | 1157010 | G | Introduction 10 |
| L3 | 2110074 | T | Introduction 10 |
| L3 | 294514 | C | Introduction 10 |
| L3 | 3221361 | C | Introduction 10 |
| L3 | 3630122 | T | Introduction 10 |
| L3 | 3636991 | A | Introduction 10 |
| L3 | 931240 | T | Introduction 10 |

Figure S1

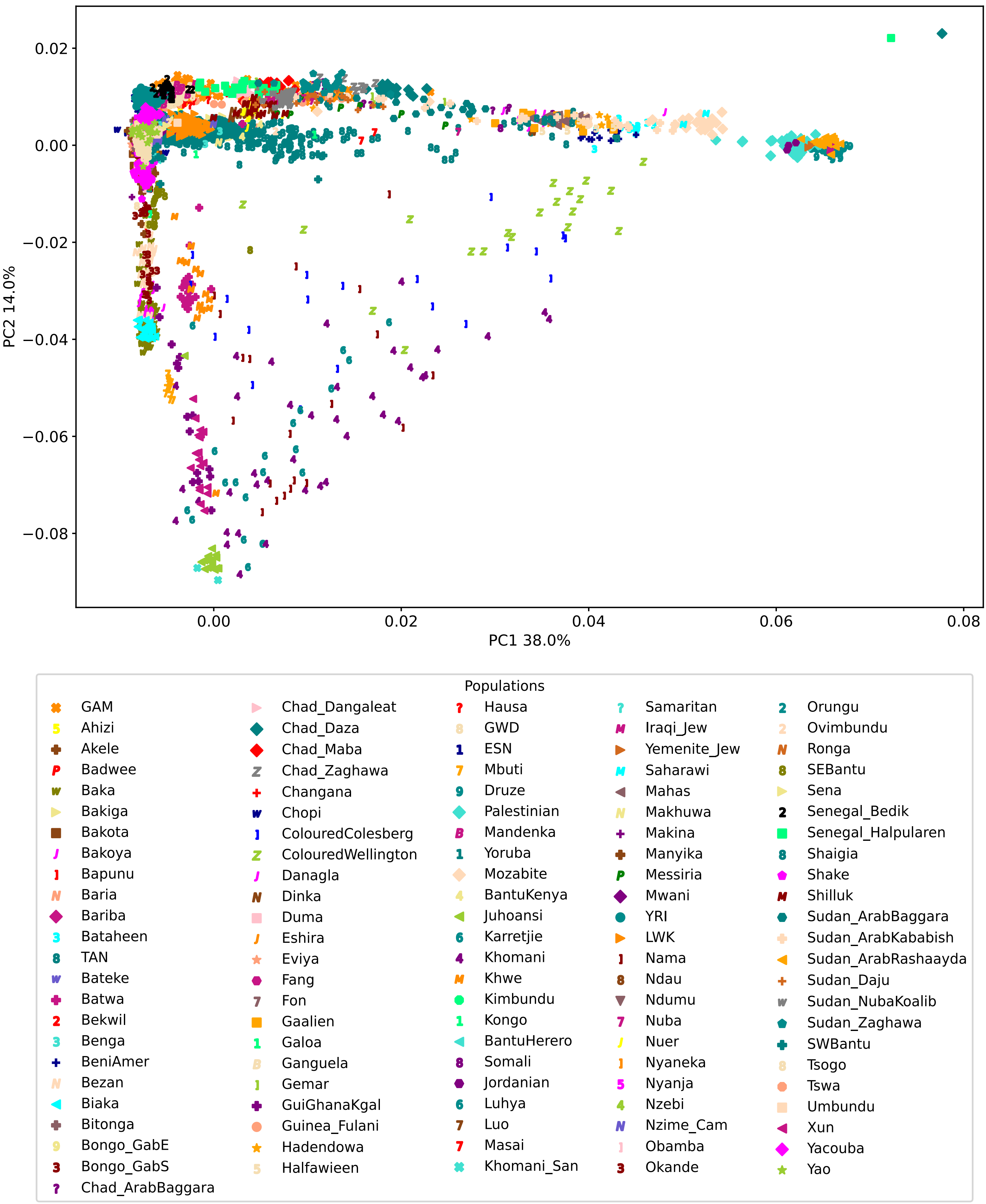
0.02
0.00
−0.02
−0.04
−0.06
−0.08
PC2 14.0%
0.00
0.02
0.04
0.06
0.08
PC1 38.0%
Populations
GAM
Ahizi
Akele
Badwee
Baka
Bakiga
Bakota
Bakoya
Bapunu
Baria
Bariba
Bataheen
TAN
Bateke
Batwa
Bekwil
Benga
BeniAmer
Bezan
Biaka
Bitonga
Bongo_GabE
Bongo_GabS
Chad_ArabBaggara
Chad_Dangaleat
Chad_Daza
Chad_Maba
Chad_Zaghawa
Changana
Chopi
ColouredColesberg
ColouredWellington
Danagla
Dinka
Duma
Eshira
Eviya
Fang
Fon
Gaalien
Galoa
Ganguela
Gemar
GuiGhanaKgal
Guinea_Fulani
Hadendowa
Halfawieen
Hausa
GWD
ESN
Mbuti
Druze
Palestinian
Mandenka
Yoruba
Mozabite
BantuKenya
Juhoansi
Karretjie
Khomani
Khwe
Kimbundu
Kongo
BantuHerero
Somali
Jordanian
Luhya
Luo
Masai
Khomani_San
Samaritan
Iraqi_Jew
Yemenite_Jew
Saharawi
Mahas
Makhuwa
Makina
Manyika
Messiria
Mwani
YRI
LWK
Nama
Ndau
Ndumu
Nuba
Nuer
Nyaneka
Nyanja
Nzebi
Nzime_Cam
Obamba
Okande
Orungu
Ovimbundu
Ronga
SEBantu
Sena
Senegal_Bedik
Senegal_Halpularen
Shaigia
Shake
Shilluk
Sudan_ArabBaggara
Sudan_ArabKababish
Sudan_ArabRashaayda
Sudan_Daju
Sudan_NubaKoalib
Sudan_Zaghawa
SWBantu
Tsogo
Tswa
Umbundu
Xun
Yacouba
Yao

Figure S2

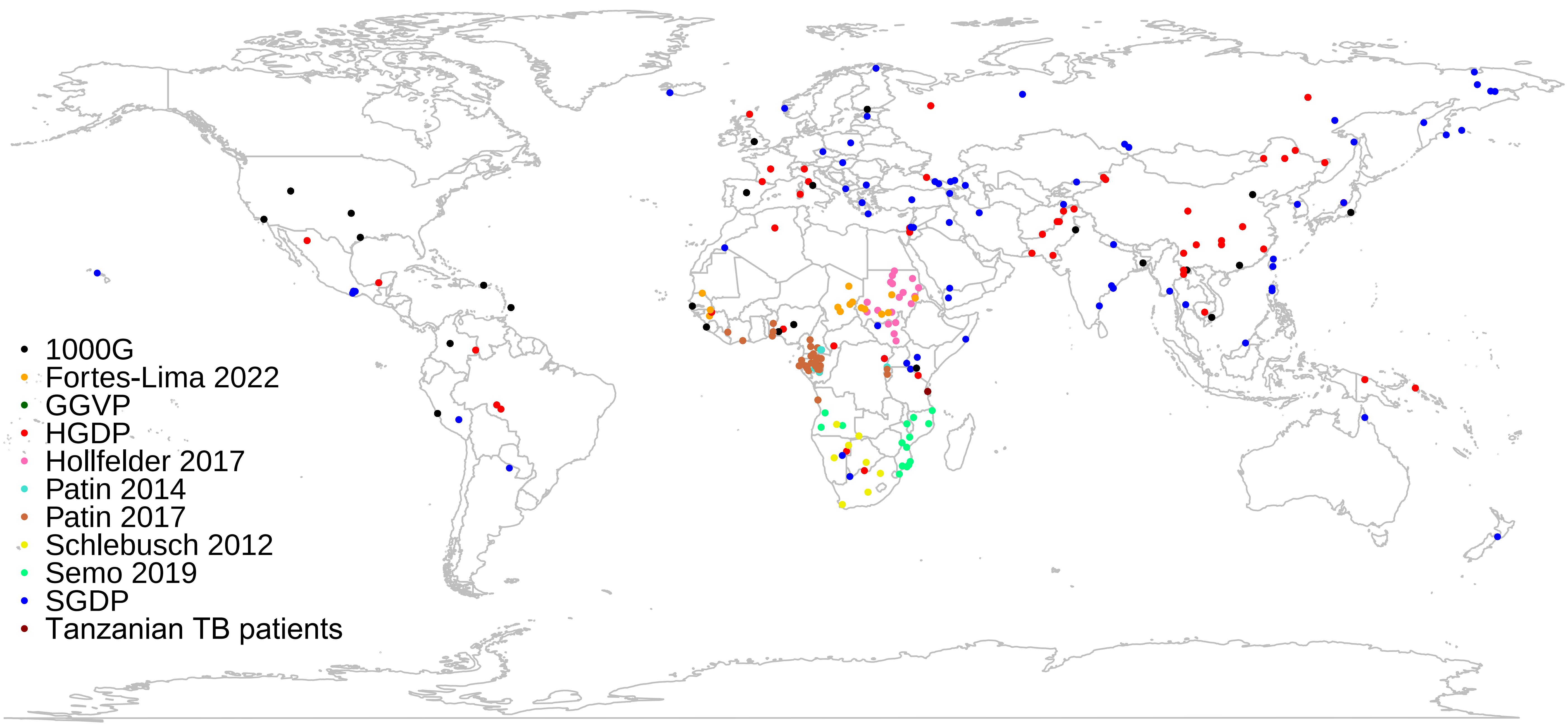

1000G
Fortes-Lima 2022
GGVP
HGDP
Hollfelder 2017
Patin 2014
Patin 2017
Schlebusch 2012
Semo 2019
SGDP
Tanzanian TB patients

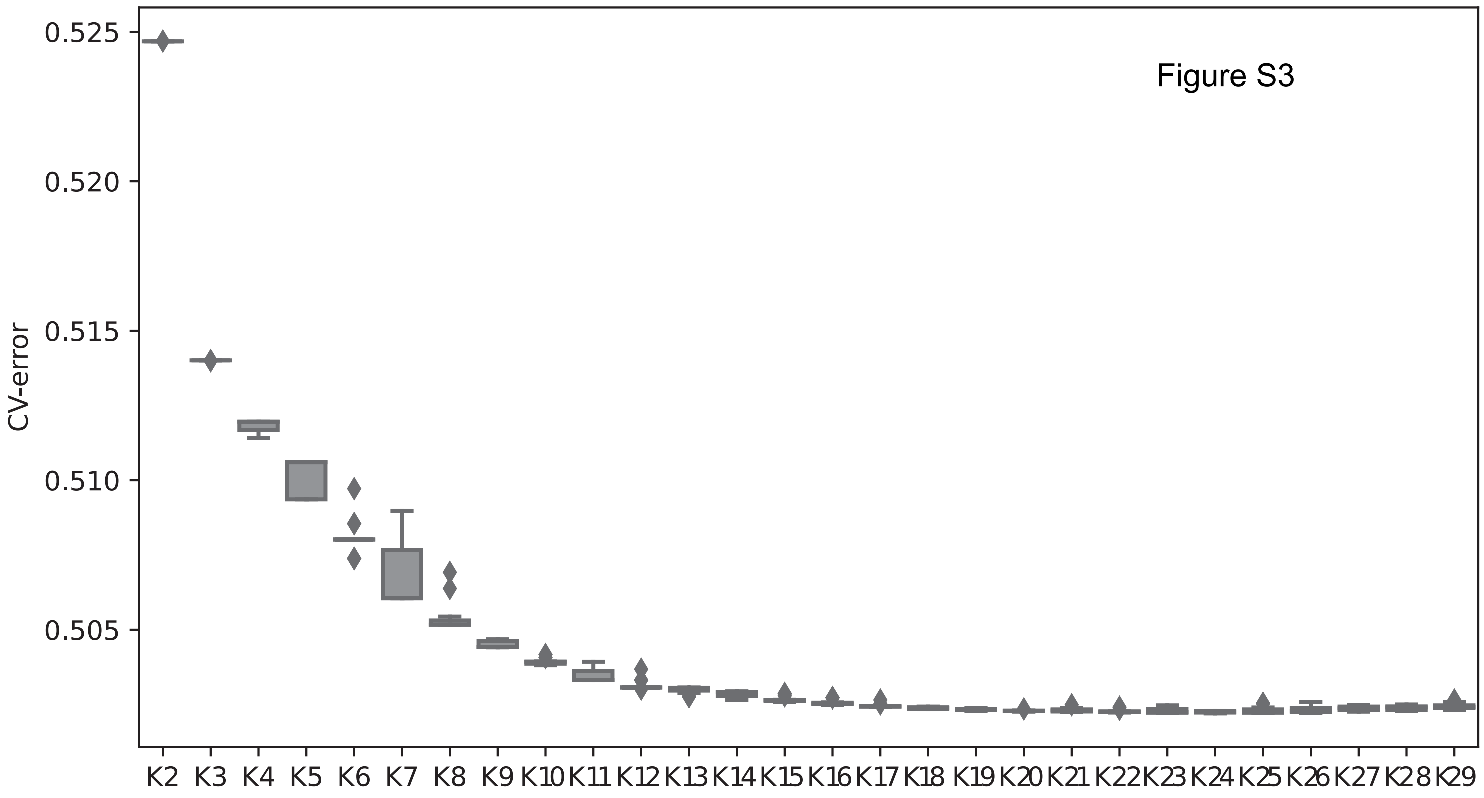

Figure S3
CV-error
0.525
0.520
0.515
0.510
0.505
K2 K3 K4 K5 K6 K7 K8 K9 K10 K11 K12 K13 K14 K15 K16 K17 K18 K19 K20 K21 K22 K23 K24 K25 K26 K27 K28 K29

Figure S4

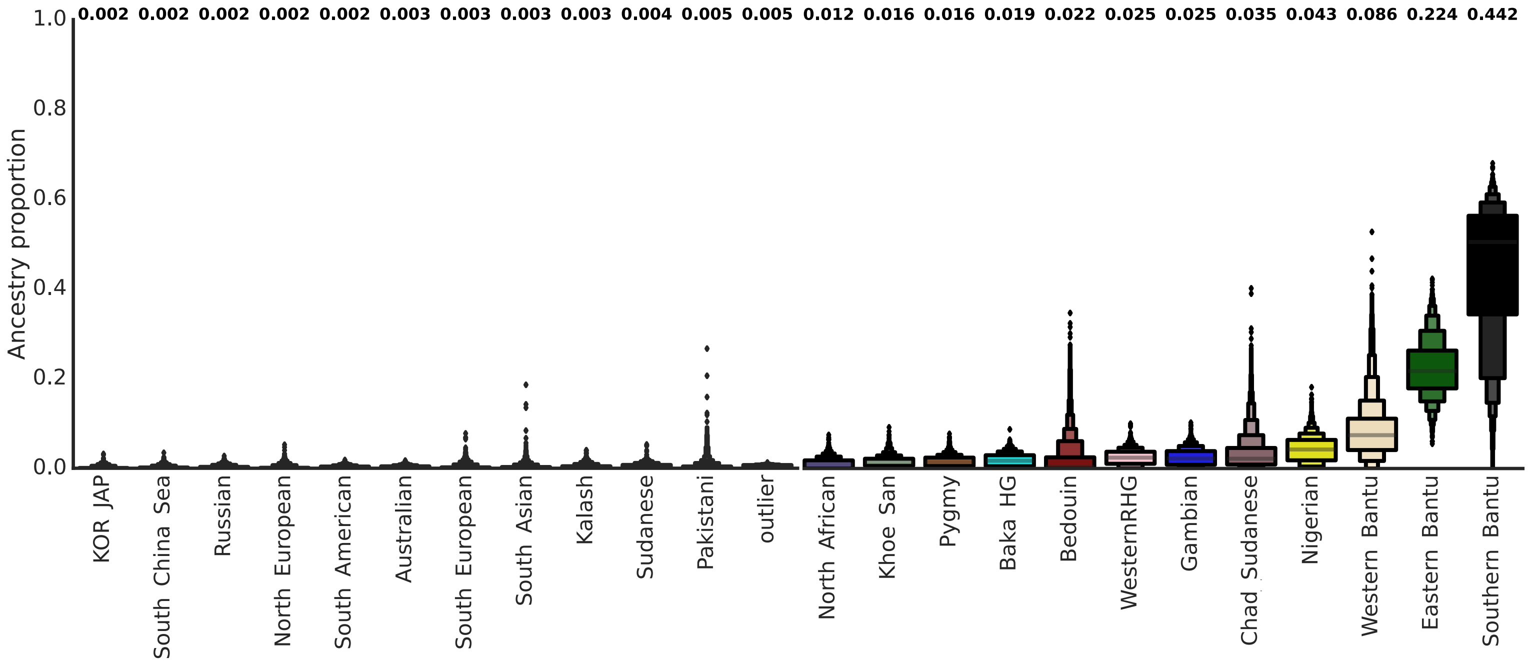

0.002 0.002 0.002 0.002 0.002 0.003 0.003 0.003 0.003 0.004 0.005 0.005 0.012 0.016 0.016 0.019 0.022 0.025 0.025 0.035 0.043 0.086 0.224 0.442
1.0
0.8
0.6
0.4
0.2
0.0
Ancestry proportion
KOR JAP
South China Sea
Russian
North European
South American
Australian
South European
South Asian
Kalash
Sudanese
Pakistani
outlier
North African
Khoe San
Pygmy
Baka HG
Bedouin
WesternRHG
Gambian
Chad Sudanese
Nigerian
Western Bantu
Eastern Bantu
Southern Bantu

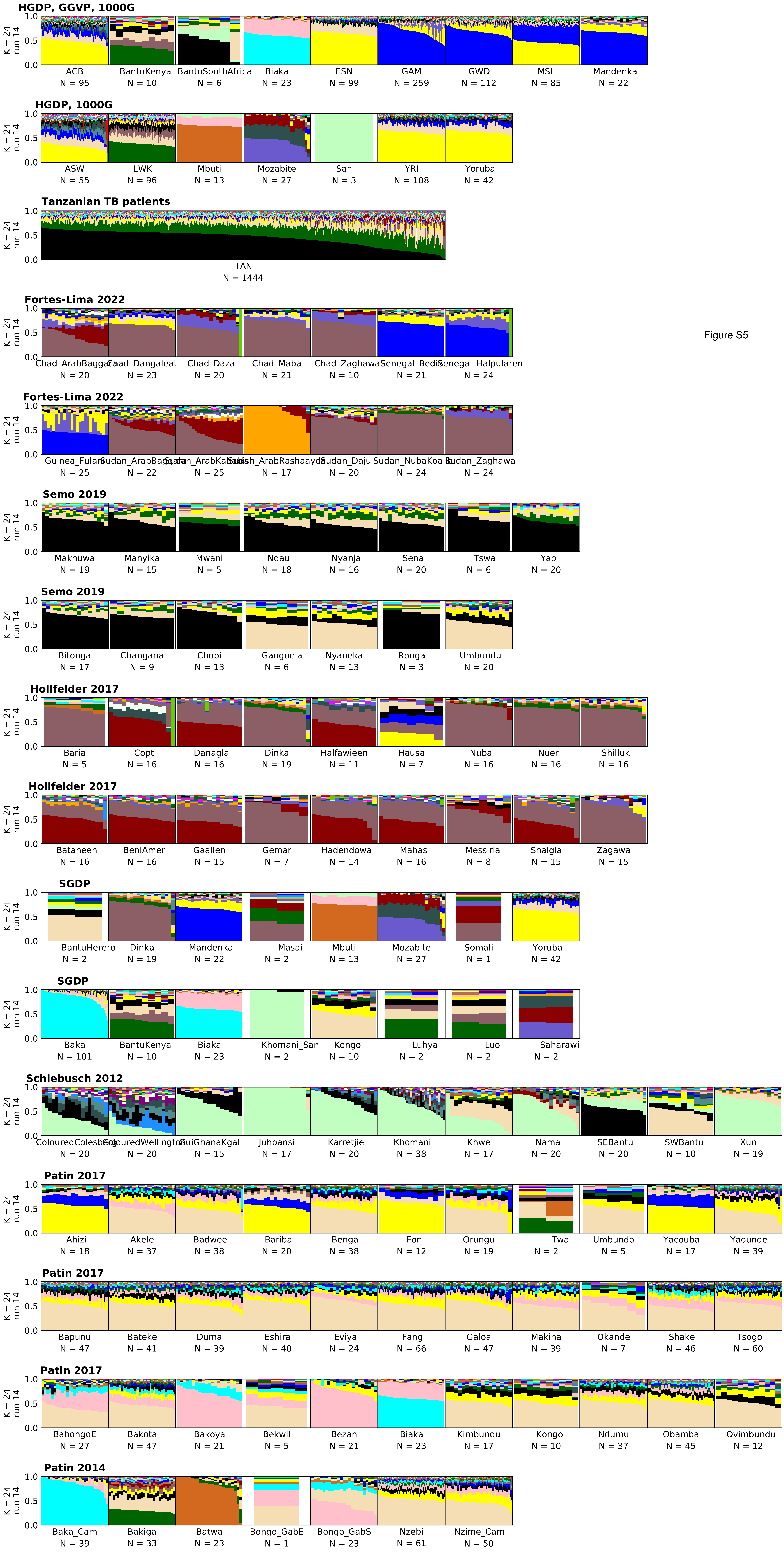
HGDP, GGVP, 1000G
K = 24 run 14
ACB N = 95
BantuKenya N = 10
BantuSouthAfrica N = 6
Biaka N = 23
ESN N = 99
GAM N = 259
GWD N = 112
MSL N = 85
Mandenka N = 22
HGDP, 1000G
ASW N = 55
LWK N = 96
Mbuti N = 13
Mozabite N = 27
San N = 3
YRI N = 108
Yoruba N = 42
Tanzanian TB patients
TAN N = 1444
Fortes-Lima 2022
Chad_ArabBaggara N = 20
Chad_Dangaleat N = 23
Chad_Daza N = 20
Chad_Maba N = 21
Chad_Zaghawa N = 10
Senegal_Bedik N = 21
Senegal_Halpularen N = 24
Fortes-Lima 2022
Guinea_Fulani N = 25
Sudan_ArabBaggara N = 22
Sudan_ArabKababish N = 25
Sudan_ArabRashaayda N = 17
Sudan_Daju N = 20
Sudan_NubaKoalib N = 24
Sudan_Zaghawa N = 24
Semo 2019
Makhuwa N = 19
Manyika N = 15
Mwani N = 5
Ndau N = 18
Nyanja N = 16
Sena N = 20
Tswa N = 6
Yao N = 20
Semo 2019
Bitonga N = 17
Changana N = 9
Chopi N = 13
Ganguela N = 6
Nyaneka N = 13
Ronga N = 3
Umbundu N = 20
Hollfelder 2017
Baria N = 5
Copt N = 16
Danagla N = 16
Dinka N = 19
Halfawieen N = 11
Hausa N = 7
Nuba N = 16
Nuer N = 16
Shilluk N = 16
Hollfelder 2017
Bataheen N = 16
BeniAmer N = 16
Gaalien N = 15
Gemar N = 7
Hadendowa N = 14
Mahas N = 16
Messiria N = 8
Shaigia N = 15
Zagawa N = 15
SGDP
BantuHerero N = 2
Dinka N = 19
Mandenka N = 22
Masai N = 2
Mbuti N = 13
Mozabite N = 27
Somali N = 1
Yoruba N = 42
SGDP
Baka N = 101
BantuKenya N = 10
Biaka N = 23
Khomani_San N = 2
Kongo N = 10
Luhya N = 2
Luo N = 2
Saharawi N = 2
Schlebusch 2012
ColouredColesberg N = 20
ColouredWellington N = 20
GuiGhanaKgal N = 15
Juhoansi N = 17
Karretjie N = 20
Khomani N = 38
Khwe N = 17
Nama N = 20
SEBantu N = 20
SWBantu N = 10
Xun N = 19
Patin 2017
Ahizi N = 18
Akele N = 37
Badwee N = 38
Bariba N = 20
Benga N = 38
Fon N = 12
Orungu N = 19
Twa N = 2
Umbundo N = 5
Yacouba N = 17
Yaounde N = 39
Patin 2017
Bapunu N = 47
Bateke N = 41
Duma N = 39
Eshira N = 40
Eviya N = 24
Fang N = 66
Galoa N = 47
Makina N = 39
Okande N = 7
Shake N = 46
Tsogo N = 60
Patin 2017
BabongoE N = 27
Bakota N = 47
Bakoya N = 21
Bekwil N = 5
Bezan N = 21
Biaka N = 23
Kimbundu N = 17
Kongo N = 10
Ndumu N = 37
Obamba N = 45
Ovimbundu N = 12
Patin 2014
Baka_Cam N = 39
Bakiga N = 33
Batwa N = 23
Bongo_GabE N = 1
Bongo_GabS N = 23
Nzebi N = 61
Nzime_Cam N = 50
Figure S5

### Figure S6

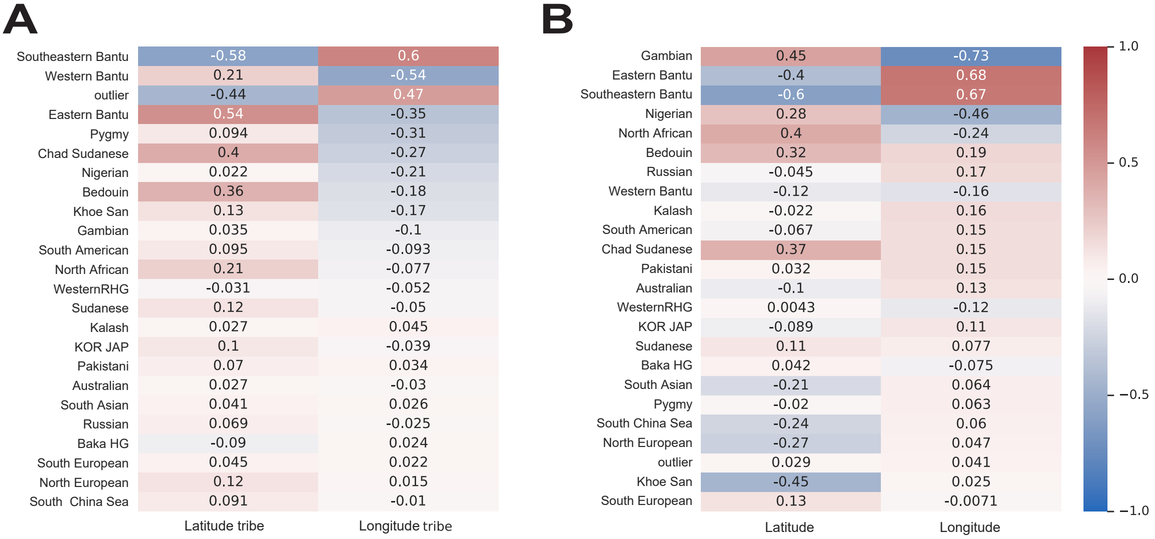
A
Southeastern Bantu -0.58 0.6
Western Bantu 0.21 -0.54
outlier -0.44 0.47
Eastern Bantu 0.54 -0.35
Pygmy 0.094 -0.31
Chad Sudanese 0.4 -0.27
Nigerian 0.022 -0.21
Bedouin 0.36 -0.18
Khoe San 0.13 -0.17
Gambian 0.035 -0.1
South American 0.095 -0.093
North African 0.21 -0.077
WesternRHG -0.031 -0.052
Sudanese 0.12 -0.05
Kalash 0.027 0.045
KOR JAP 0.1 -0.039
Pakistani 0.07 0.034
Australian 0.027 -0.03
South Asian 0.041 0.026
Russian 0.069 -0.025
Baka HG -0.09 0.024
South European 0.045 0.022
North European 0.12 0.015
South China Sea 0.091 -0.01
Latitude tribe
Longitude tribe
B
Gambian 0.45 -0.73
Eastern Bantu -0.4 0.68
Southeastern Bantu -0.6 0.67
Nigerian 0.28 -0.46
North African 0.4 -0.24
Bedouin 0.32 0.19
Russian -0.045 0.17
Western Bantu -0.12 -0.16
Kalash -0.022 0.16
South American -0.067 0.15
Chad Sudanese 0.37 0.15
Pakistani 0.032 0.15
Australian -0.1 0.13
WesternRHG 0.0043 -0.12
KOR JAP -0.089 0.11
Sudanese 0.11 0.077
Baka HG 0.042 -0.075
South Asian -0.21 0.064
Pygmy -0.02 0.063
South China Sea -0.24 0.06
North European -0.27 0.047
outlier 0.029 0.041
Khoe San -0.45 0.025
South European 0.13 -0.0071
Latitude
Longitude
1.0
0.5
0.0
−0.5
−1.0

### Figure S7

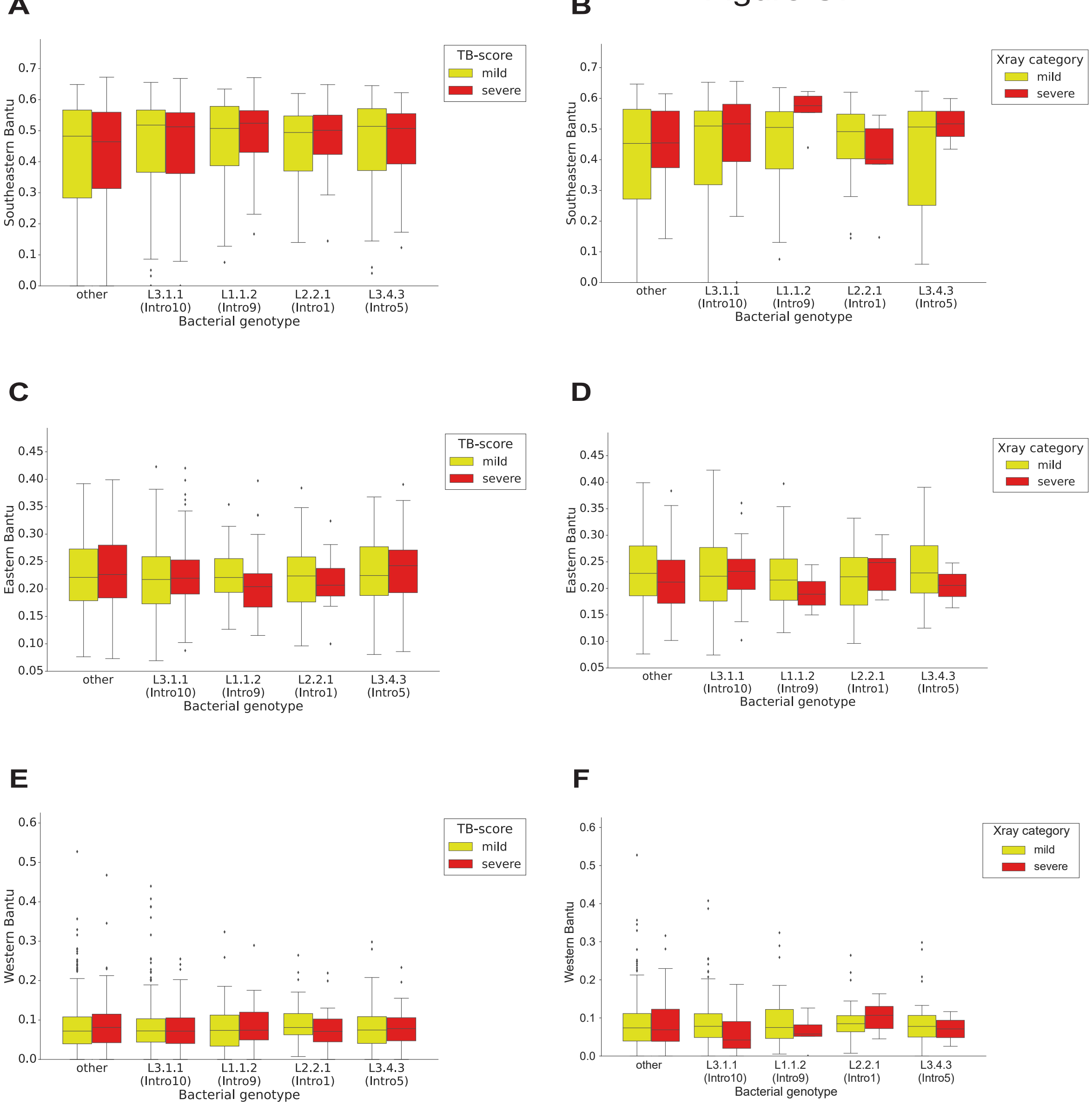
A
B
C
D
E
F
TB-score
Xray category
mild
severe
Southeastern Bantu
Eastern Bantu
Western Bantu
other
L3.1.1
(Intro10)
L1.1.2
(Intro9)
L2.2.1
(Intro1)
L3.4.3
(Intro5)
Bacterial genotype
